# Piezo1 potentially mediates inflammation in balloon-inflated rat brain and its bidirectional mechanosensitivity

**DOI:** 10.1101/2020.07.16.207589

**Authors:** Yichi Zhang, Gang Wang, Minjie Xie, Lifei Lian, Yongjie Xiong, Feng Xu, Guo Li, Zhouping Tang, Furong Wang, Suiqiang Zhu

## Abstract

Brain injury after intracerebral hemorrhage is extremely complicated, and the exact mechanism remains puzzling. Piezo1, a novel mammalian mechanosensitive ion channel, has been identified to play important roles in several pathologic and physiologic procedures that involve cellular mechanotransduction. However, the role of Piezo1 in hematoma compression after intracerebral hemorrhage is still unclear. In the present study, we established a balloon-inflated rat brain model mimicking the pure mechanical compression of a hematoma and detected balloon compression in the basal ganglia region of the brain, resulting in abnormal behaviors and a significant increase in the expression of Piezo1 and proinflammatory cytokines. These effects were reversed by GsMTx4, an antagonist of Piezo1. Additionally, the balloon deflation time affected behavioral function and the levels of Piezo1 and proinflammatory cytokines. These results establish the first *in vivo* evidence for the role of Piezo1 in blood-brain neuroinflammation after hematoma compression. Piezo1 may therefore be a potential therapeutic target for the treatment of intracerebral hemorrhage.

## Introduction

Intracerebral hemorrhage (ICH) is the second major subtype of stroke with high mortality (Fazekas *et al*., 2018).The brain injury observed after ICH is usually divided into “primary” and “secondary” injury. For the former, the compression of the circumjacent brain tissue by the hematoma, also known as the mass effect, is a crucial factor. The mass effect may induce brain parenchyma dislocation, midline shifting, and even cerebral hernia if the hematoma volume is sufficiently large (Keep *et al*., 2012). For secondary injury, the mechanism is more complicated. There are many damaging factors associated with hematoma lysis, such as erythrocyte disruption, thrombin, and the release of hemoglobin, heme, and iron ions (Zhou *et al*., 2014). Peripheral tissue compression occurs immediately upon formation of the hematoma, and this mechanical stimulus induces damage directly. However, our knowledge of the mass effect is incomplete. The effects on the cells and their response to the mechanical stimulus remain elusive.

Mechanosensitive ion channels (MSCs) are channels that sense mechanical force stimuli. However, few MSCs have been identified to date (Chalfie, 2009). DEG/ENaC ion channels were first detected to be related to light touch sensation in *Caenorhabditis elegans* (Driscoll and Chalfie, 1991; O’Hagan *et al*., 2005). In the past decade, transient receptor potential (TRP) channels were also considered to be candidate MSCs, with evidence that TRPN (one subtype of TRP) was expressed in *Drosophila melanogaster* and *Caenorhabditis elegans* (Christensen and Corey, 2007). Double-pore K^+^ (K2P) channels were the first channels found to be sensitive to mechanical force in mammals (Fink *et al*., 1996), playing key roles in modulating cytomembrane potentials (Enyedi and Czirják, 2010; Brohawn, 2015).

Recently, the identification of the Piezo family furthered our knowledge of MSCs in mammals (Coste *et al*., 2010). Briefly, Piezos are a unique class of nonselective cation channels, with the large size distinguishing them from any other known channels. Moreover, these channels are highly evolutionarily conserved across different species (Volkers *et al*., 2015). For mammals there are two homologues Piezo1 and Piezo2, which are expressed in different tissues (Xu, 2016). When purified and reconstituted in lipid bilayers, Piezos present channel activity, suggesting that these proteins are real mechanically activated channels (Coste *et al*., 2012; Coste *et al*., 2015). Piezo2 was found to be expressed in a subgroup of mouse dorsal root ganglion (DRG) neurons that are located in mammalian skin and folliculus pili and form the Merkel-neurite complex, suggesting that Piezo2 is the primary sensory channel for light touch in mammals (Maksimovic *et al*., 2014; Ikeda *et al*., 2014; Bron *et al*., 2014; Ranade *et al*., 2014*a*). Piezo1 plays pivotal roles in the development of vascular systems in mice through its sensation of blood shearing force (Ranade *et al*., 2014*b*; Li *et al*., 2014). Piezo1 mediates the cation influx of erythrocytes under compression and is involved in the modulation of cellular volume and function (Faucherre *et al*., 2014; Cahalan *et al*., 2015). In central nervous system (CNS), Piezo1 senses local surroundings, hence affecting downstream cellular signaling, differentiation and motility (Pathak *et al*., 2014; Hung *et al*., 2016), and mediates the interaction between astrocytes and neurons (Blumenthal et al., 2014). It has been reported that Hib (an equivalent of Piezo1) is over-expressed in rat astrocytes associated with senile plaques (Satoh et al., 2006). With specific knockout of Piezo1 in smooth muscle cells, mice survive but present a functional deficiency of vascular remolding under hypertension (Retailleau *et al*., 2015). In addition, Piezo1 senses fluid shearing force and mediates the mechanically activated current in bladder urothelial cells and renal endothelial cells (Martins *et al*., 2016).

To the best of our knowledge, the expression and roles of Piezos in brain injury after ICH remain elusive. As novel mammalian mechanosensitive channels, Piezos are most likely involved in the response of neural cells to compression resulting from a hematoma. In the present study, a balloon-inflated rat model was established to mimic the mechanical force in the basal ganglia associated with a hematoma without the influence of blood components, thus allowing us to focus on the pure mass effect of hematoma. We examined the expression level of Piezo1, the effects of Piezo1 on proinflammatory factors, including interleukin (IL)-1β, IL-6, tumor necrosis factor (TNF-α), and the behavioral performance of rats after balloon inflation brain injury.

## Materials and Methods

### Ethics statement

The entire experiment was authorized by the Laboratory Animal Welfare and Ethics Committee of Tongji Hospital, Huazhong University of Science and Technology. We minimized the number of rats used and the pain they suffered to the best of our ability.

### Animal model

The animal model of this study was established on male Sprague-Dawley rats 10 weeks old and 300-400 g in weight. Animals were maintained in a specific pathogen-free (SPF) environment with constant temperature and humidity and free access to food and water for at least three days to become familiar with the environment. For the uniformity, the operation is always carried out at 10:00 in the morning. Animals were anesthetized by an intraperitoneal injection of 6% chloral hydrate (the most efficient and safe anesthesia for rats) and then positioned on a standard stereotaxic apparatus (*RWD life science, China*) to present a flat skull. The coordinates for the right basal ganglia were 2.12 mm posterior to bregma, 3.5 mm rightward of the midline, and 6 mm deep from the surface of the duramater (Paxinos and Watson, 1996). The rats were divided randomly (randomization procedure was achieved at http://www.randomizer.org/) into a sham group (pseudo-operation control), a model group (mechanical compression injury), and intervention groups (GsMTx4 administration or different deflation times). Each group contained six rats. For the sham group, the balloon catheter (*PTCA, Boston Scientific, USA*) was embedded but not inflated. For the model group, the balloon was embedded, inflated to a pressure of 0.5 ATM with a volume of approximately 25 μL, and maintained for 30 min; the balloon was then deflated and displaced in one minute. In the GsMTx4 group, GsMTx4 (*3* μ*M, ab141871, Abcam, UK*), an antagonist of Piezo1, was applied before the balloon insertion. In the groups with different deflation times, the balloon was deflated in either ten seconds or five minutes, while the preceding operations were all the same as those in the model group. After the operation, the skin was sewn, and animals were returned to the SPF environment to wake up from anesthesia. During the above procedures, the body temperature of the animals was maintained at approximately 37 °C. All operations were performed under aseptic conditions. The neural function of the animals was assessed using the Garcia scale over three consecutive days, and the assessment was performed by a researcher who was blinded to the groups. During the whole experiment, totally 26 rats were wasted owing to various reasons, such as unexpected death and absent of syndromes. The graphical time line is presented below:

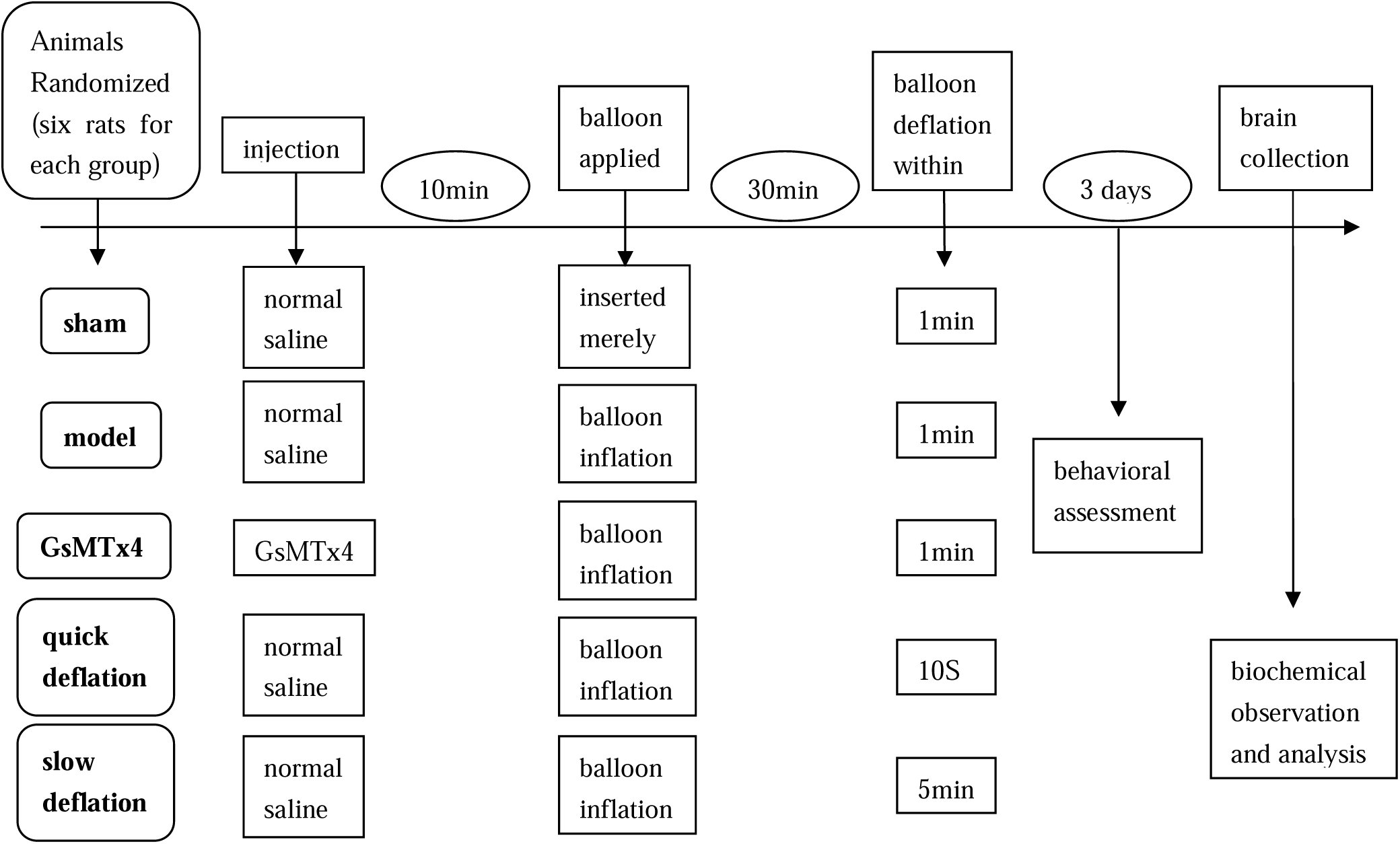

### Garcia scale assessment

After the operation, the animals were assessed functionally using the Garcia scale, which contains six terms and a total score of 18, with a higher score indicating better neurological function. For the behavioral observation, spontaneous activity, symmetry of movements, forelimb outstretching, wire cage wall climbing, reaction to touch on either side of the trunk and response to vibrissae touch were assessed. The detailed criteria can be seen in the report by Garcia *et al*. (Garcia *et al*., 1995). At the 72 h point after operation, rats were over-anesthetized to death for the preparation of following experiments.

### Immunofluorescence

Brains were collected and stored at −80 °C, and 10-µm-thick frozen sections were prepared at -20 °C (*Leica CM1950, Germany*). Sections were stored at -80 °C and used for immunofluorescence. After the sections were fixed in precooledacetone ethanol (4 °C, 1:1) for 20 min, they were treated with 3% H_2_O_2_ for 10 min to inactivate endogenous peroxidases and then closed in 5% bovine serum albumin (BSA; *10735078001, Roche, Switzerland*) for 20 min. Sections were then covered with the primary antibody anti-FAM38A (*1:200, ab128245, Abcam, UK*) at 4 °C overnight. After primary antibody treatment, the sections were incubated with a FITC-labeled second antibody (*1:200, AS-1112, Aspen, China*) at 37 °C for 50 min and then stained with 4’,6-diamidino-2-phenylindole (DAPI; *AS-1075, Aspen, China*) at room temperature for 5 min away from light. After the sections were treated with anti-quenching agent (*AS-1089, Aspen, China*), they were observed and photographed on an Olympus system consisting of a microscope and a digital camera (Olympus BX53; Olympus Fluoview FV1200; Olympus CellSens V1.6).

### Western blotting

The level of the target protein Piezo1 was assessed by Western blotting. At 72h after stereotaxic operation, the rats were anesthetized by 6% chloral hydrateintraperitoneal injection and immediately decapitated for collection of the right brain hemispheres, which were frozen in liquid nitrogen for 30 min and stored at -80 °C. Brain samples were thawed at room temperature and ground in ice-cold Tris-buffered saline (TBS) containing a Protease Inhibitor Cocktail Tablet (*Roche, Switzerland*), with a final dilution of 1:100. The homogenates were centrifuged for 30 min at 13,000 g at 4 °C, and the supernatants were collected. The concentration of Piezo1 was determined using a BCA Protein Assay Kit (*AS1086, Aspen, China*). For each sample, 40 μg of total protein was separated on a 10% SDS-PAGE gel and transferred onto a nitrocellulose membrane, which was blocked in 5% fat-free milk for 30 min at 20 °C and incubated overnight at 4 °C with the primary antibody anti-FAM38A (*1:2000, ab128245, Abcam, UK*). Then, the samples were incubated with the secondary antibody (*HRP-goat anti-rabbit, 1:10000, AS1107, Aspen, China*) for 1 h at 37 °C. After treatment with enhanced chemiluminescence reagents (*AS1059, Aspen, China*), the membrane was scanned (*LiDE110, Canon, Japan*), and the optical density was determined using ImageJ software (*v1*.*46; National Institutes of Health, USA*).

### Enzyme-linked immunosorbent assay (ELISA) kits

Brain samples were homogenized with ice-cold PBS and centrifuged at 4 °C, 12000 rps for 20 min. Then, the supernatants were harvested and stored at-80 °C. The levels of the proinflammatory factors IL-1β, IL-6, and TNF-α were measured by commercial ELISA kits (*QuantiCyto, Neobioscience Company of Biotechnology, China*) following the manufacturer’s instructions.

### Statistical analysis

The quantitative data are presented as the means ± standard error of the mean (SEM). Statistical analysis was carried out using SPSS Statistics (*version 22*.*0, IBM, USA*). The two-tailed Student’s t test was performed for comparisons between two groups. Mauchly’s test of sphericity and multiway ANOVA followed by the least significant difference test were used for comparisons of multiple groups. A significant difference was accepted as a P value<0.05 (*GraphPad Prism version 5*.*00*).

### Data availability

The data will be shared on request of a qualified investigator for any reasonable noncommercial research purposes within the limits of all authors’ consent.

## Results

### 1. Balloon inflation formed a mass effect focused in the basal ganglia and effectively induced neurologicalfunctional deficiency mimicking ICH injury in a rat model

After the balloon inflation operation, rats woke up from anesthesia approximately three hours later and presented hemiplegic symptoms, including deficiencies in walking, climbing, balancing and touch response. Immunofluorescence revealed an obvious cavity in the basal ganglia region of DAPI-stained brain sections **(Fig. 1A, B)**. Assessment using the Garcia scale over three consecutive days revealed that those in the model group displayed prominently worse behaviors than those in the sham group (P<0.05) **(Fig. 1C)**.

**Figure 1.**
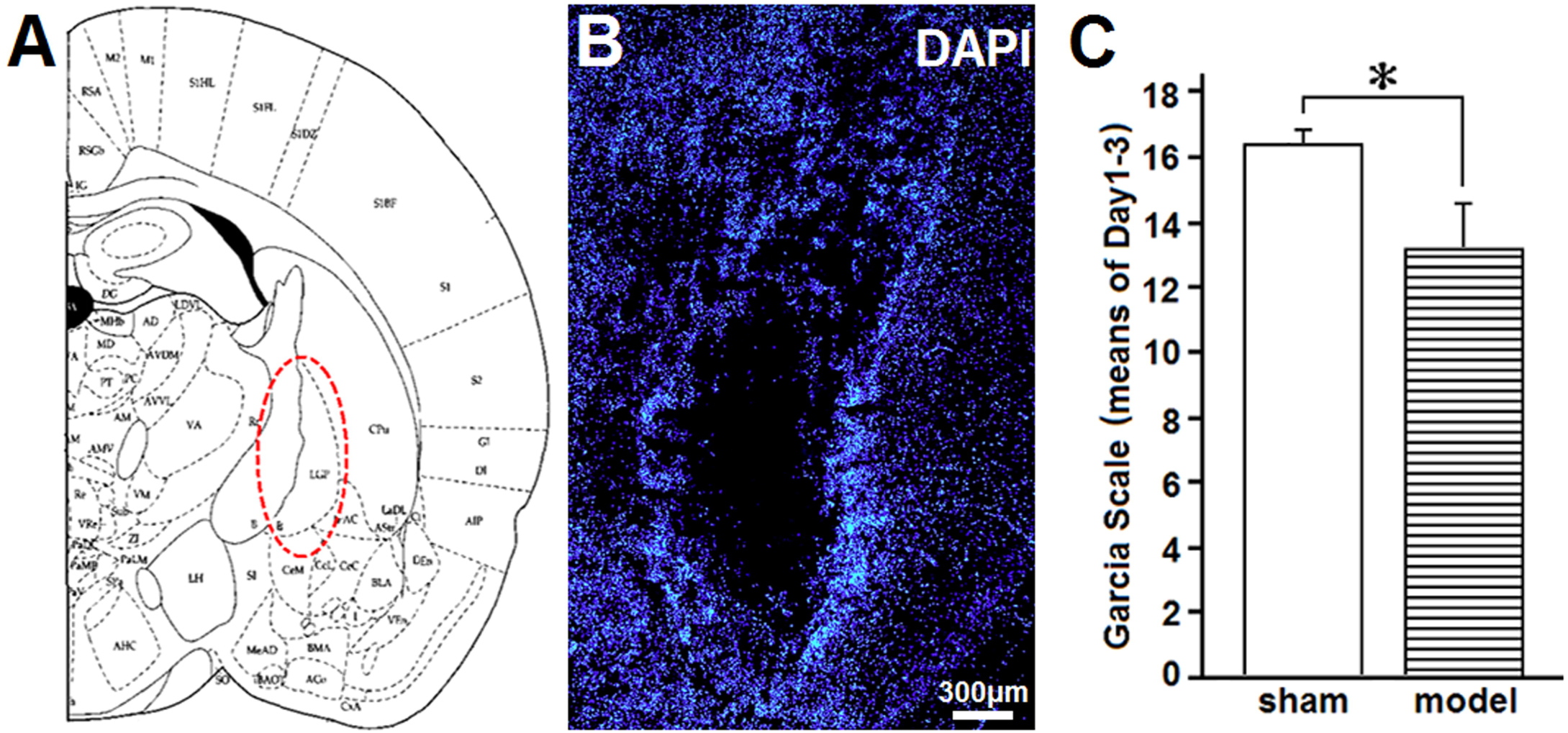
The epicenter of the balloon inflation brain injury and comparison of the behavioral outcome between the sham and model groups. A: A sketch of the right hemisphere of the rat brain based on Paxinos and Watson (Paxinos and Watson, 1996). The red dashed ellipse shows the approximate region injured by balloon inflation, mainly including the capsula interna. B: A DAPI-stained brain section of a model rat. A cavity can be seen in the expected epicenter of the injury. The bar indicates 300 μm. C: The Garcia scale was applied to evaluate the functional injury after modeling operation; “sham” means the balloon was inserted but not inflated, and “model” means that the balloon was inserted and inflated for 30 min. Multiway ANOVA followed by the least significant difference test was used for statistical analysis. The asterisk indicates a significant difference (P<0.05). (n=6 for the rats number of each group)

### 2. Expression of Piezo1 protein was detected around the balloon inflation epicenter and was elevated in the presence of compression

In the sham group, Piezo1 protein immunofluorescence was detected along the edge of the tissue displaced by the insertion of the uninflated balloon. The fluorescence intensity was higher in the model group than in the sham group and presented an elliptical shape surrounding the region in which the balloon was inserted and inflated (**Fig. 2A)**. Quantitative analysis of 287-kDa Western blot bands showed that Piezo1 protein expression was significantly higher in the model group than in the sham group (P<0.05) **(Fig. 2B, C)**.

**Figure 2.**
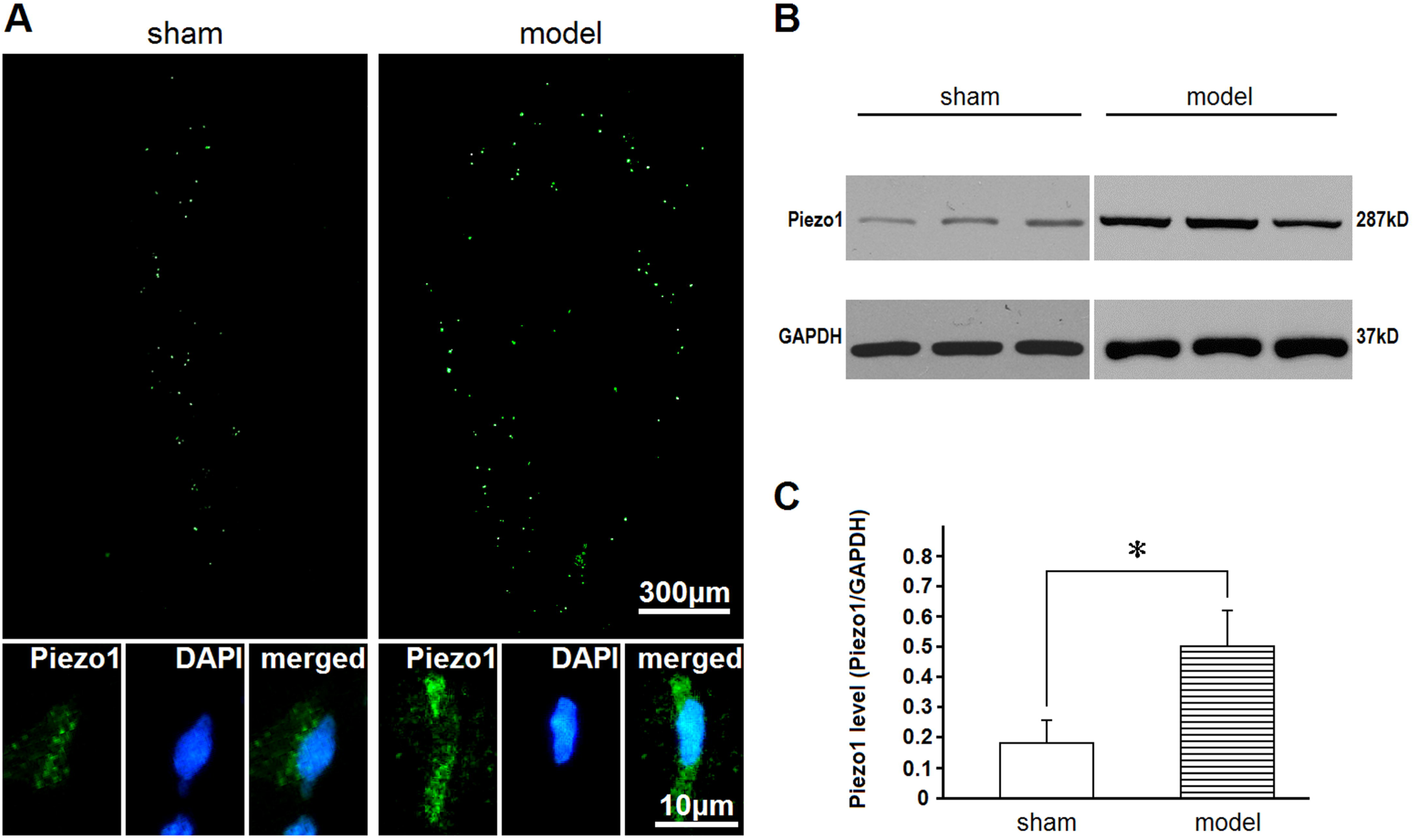
Comparison of Piezo1 expression between the sham and model groups. A: Upper channels show the immunofluorescence of Piezo1 around the balloon-shaped foci, and bottom channels show the merged magnifications identifying Piezo1 protein. DAPI=4’, 6-diamidino-2-phenylindole. B: Western blotting qualitative observation of Piezo1. Glyceraldehyde-3-phosphate dehydrogenase (GAPDH) was used as an internal control. C: Quantitative analysis of Piezo1 expression using Western blotting. Student’s t test was used for statistical analysis. The asterisk indicates a significant difference (P<0.05). (n=3 for the rats number of each group) “sham” means the balloon was inserted but not inflated. “model” means that the balloon was inserted and inflated for 30 min.

### 3. Proinflammatory cytokines expression was increased in balloon-inflated brains from the model group

The rats were euthanized on the 3rd day after the operation, and brains were collected. ELISA kits were then used to measure proinflammatory cytokine expression in the brain. As typical classic proinflammatory cytokines, IL-1β, IL-6 and TNF-α were selected for examination. The expression of these cytokines was significantly higher in the rats in the model group than in those in the sham group (P<0.05) **(Fig. 3)**.

**Figure 3.**
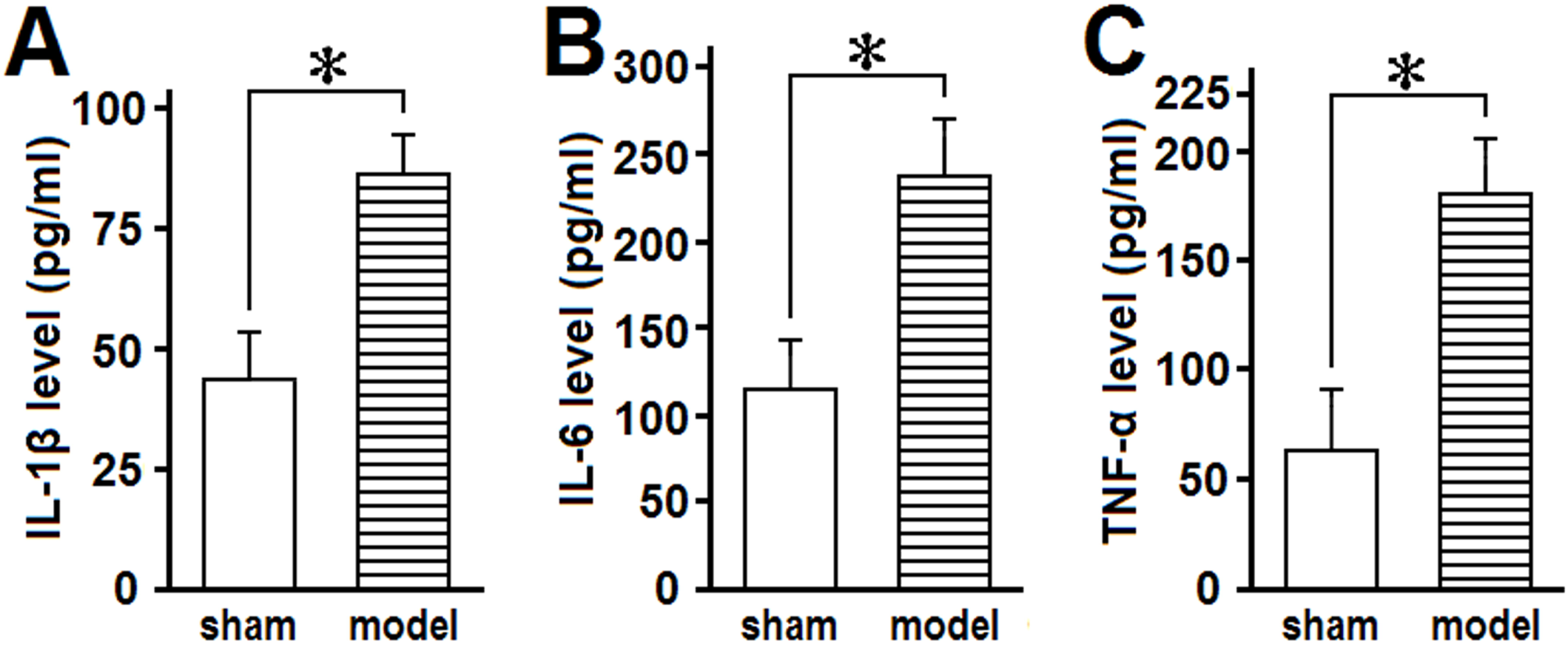
Comparison of levels of proinflammatory cytokines between the sham and model groups. Classic proinflammatory cytokines IL-1β, IL-6 and TNF-α were quantitatively analyzed and compared between the sham and model groups. Student’s t test was used for statistical analysis. Asterisks indicate significant differences (P<0.05). (n=3 for the rats number of each group) “sham” means the balloon was inserted but not inflated. “model” means that the balloon was inserted and inflated for 30 min.

### 4. GsMTx4 improved the behavioral outcome and reduced proinflammatory cytokine expression in balloon-inflated brains

Rats in the GsMTx4 group were treated with GsMTx4 (3 μM) before balloon inflation, and the potential effects were observed. The behavioral outcome of the GsMTx4 group assessed by the Garcia scale was significantly better than that of the model group (P<0.05) (**Fig. 4A**). Furthermore, in the GsMTx4 group, the levels of proinflammatory cytokines were remarkably lower than those in the model group (P<0.05) (**Fig. 4B-D**).

**Figure 4.**
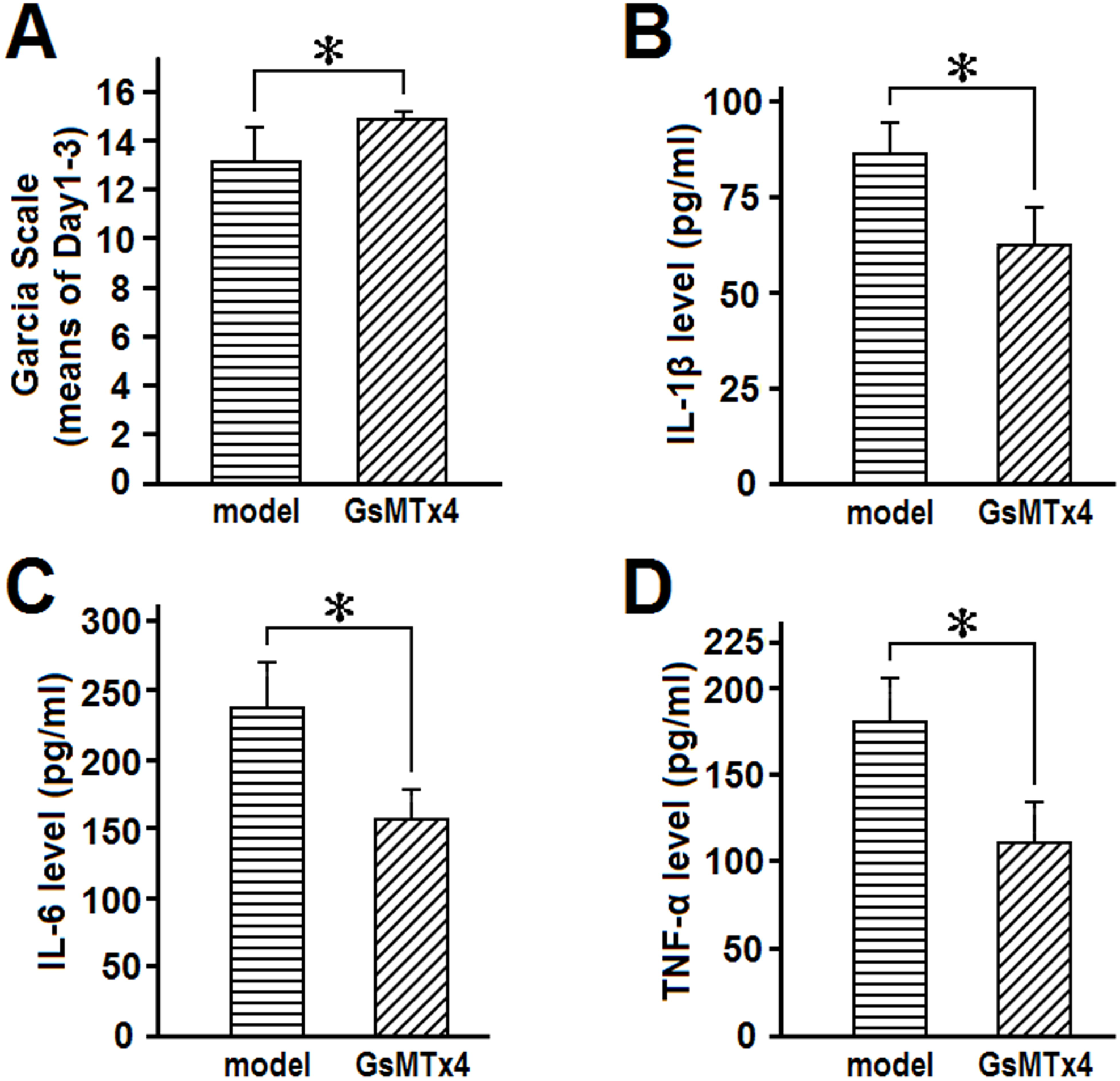
Effects of GsMTx4 on behavioral outcome and proinflammatory cytokine expression. A: The Garcia scale was applied to evaluate the behavioral outcome for three consecutive days after stereotaxic operation. B-D: The levels of classic proinflammatory cytokines IL-1β, IL-6 and TNF-α were quantitatively analyzed and compared between the model and GsMTx4 groups. Multiway ANOVA followed by the least significant difference test and Student’s T test were used for statistical analysis. Asterisks indicate significant differences (P<0.05). (n=6 for the rats number of each group) “model” means the balloon was inserted and inflated for 30 min, and “GsMTx4” means application of GsMTx4 prior to “model” operation.

### 5. The behavioral outcome was dependent on balloon deflation time

In addition to the duration of tissue compression (data not shown), the deflation time also affected the behavioral changes caused by balloon inflation-induced brain tissue compression. After balloons were inflated for 30 min, we deflated the balloons over 10 s, 1 min or 5 min and assessed the functional behavior of the rats using the Garcia scale every day for three days. Interestingly, compared to the model group, in which balloon deflation time was 1 min, the group with a deflation time of 10 s demonstrated a much lower score on the Garcia scale (P<0.05). Moreover, the Garcia scale score in the group with the deflation time prolonged to 5 min was slightly but not significantly lower than that in the group with a 1-min deflation time (P>0.05) **(Fig. 5)**.

**Figure 5.**
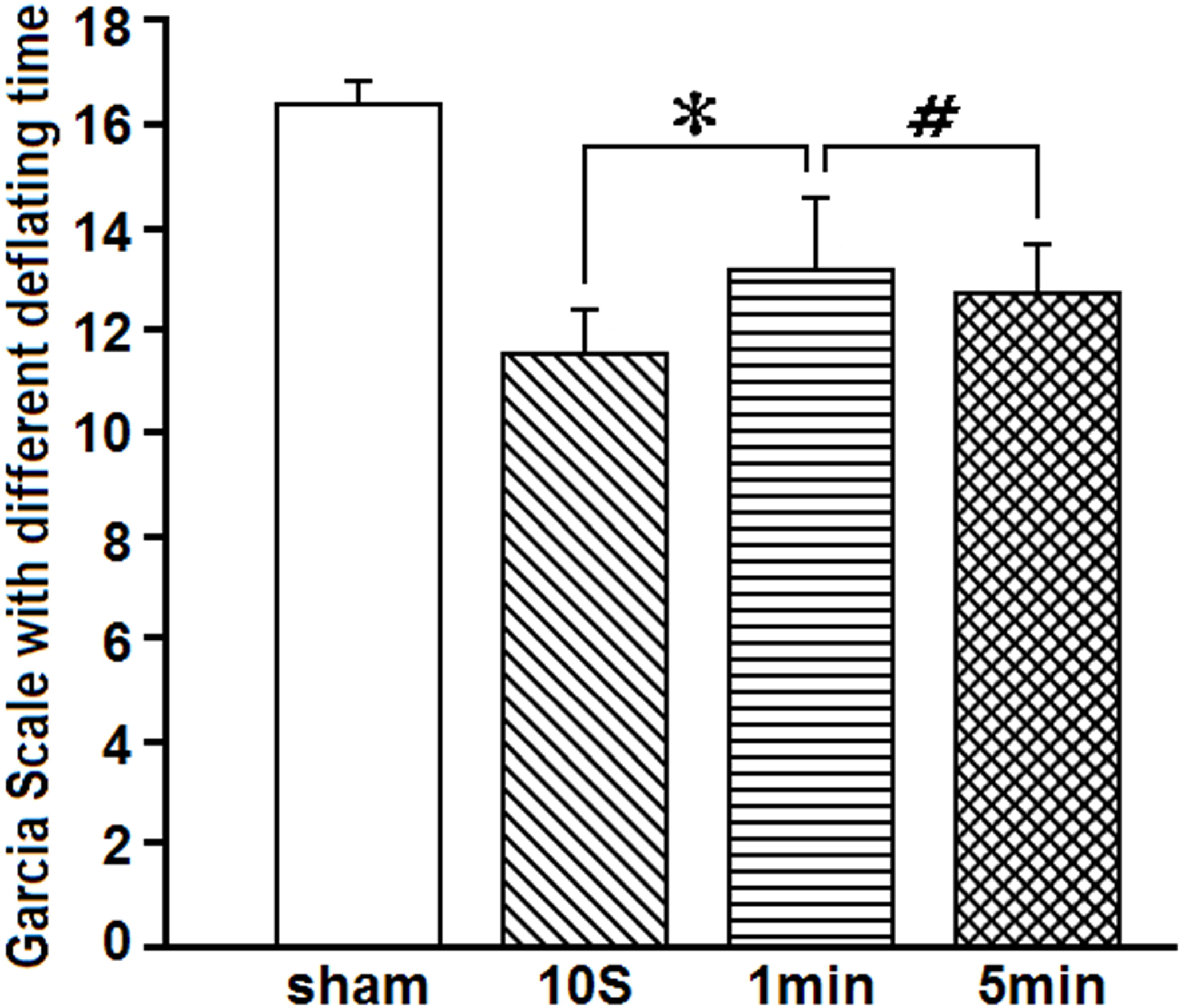
Comparison of behavioral outcomes among groups with different balloon deflation times. The Garcia scale was applied to compare the behavior outcome of groups with different balloon deflation times. “10 s”, “1 min”, and “5 min” refers to the deflation time after the balloon was inflated for 30 min. Multiway ANOVA followed by the least significant difference test was used for statistical analysis. Asterisks indicate significant differences (P<0.05), and hash indicates no significant differences (P>0.05). (n=6 for the rats number of each group)

### 6. The effects of balloon deflation time on Piezo1 expression

Furthermore, the expression of Piezo1 was examined after the rats were exposed different balloon deflation times. Quantitative analysis of the balloon-shaped distribution of Piezo1 immunofluorescence in brain sections **(Fig. 6A-D)** indicated that Piezo1 expression was higher in the group with a 10-s deflation time than in the group with a 1-min deflation time (P<0.05). However, Piezo1 expression levels in the groups with a 5-min and 1-min deflation time were not significantly different (P>0.05) **(Fig. 6E, F)**.

**Figure 6.**
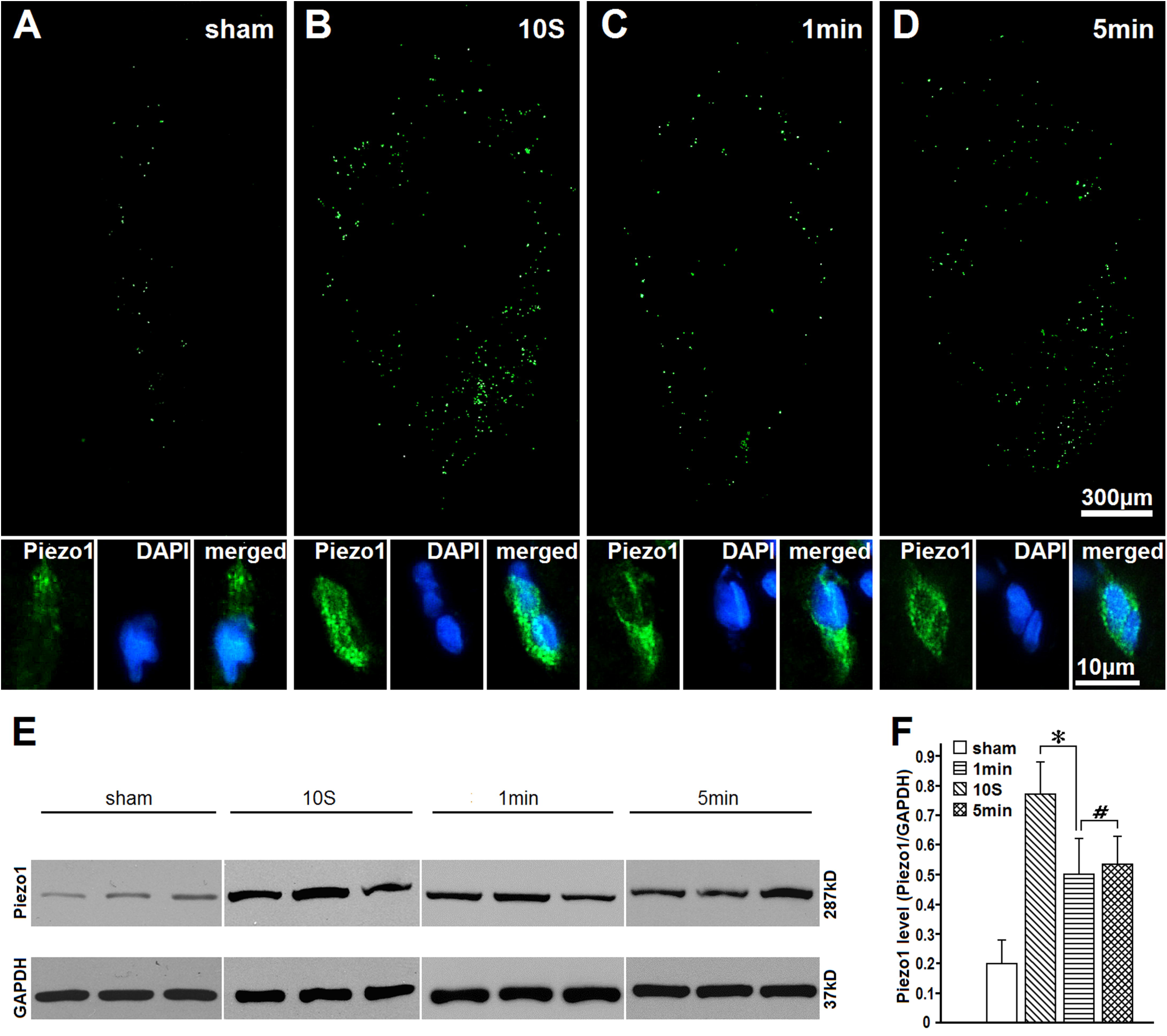
Comparison of Piezo1 expression among groups with different balloon deflation times. (A-D): Upper channels show Piezo1 immunofluorescence around the balloon-shaped foci, and bottom channels show magnified images of Piezo1 and DAPI expression. DAPI=4’, 6-diamidino-2-phenylindole. E: Western blotting of Piezo1. GAPDH was used as an internal control. F: Quantitative analysis of Piezo1 expression using Western blotting. “10 s”, “1 min”, and “5 min” refers to the deflation time after the balloon was inflated for 30 min. Student’s t test was used for statistical analysis. Asterisks indicate significant differences (P<0.05), and hash indicates no prominent differences (P>0.05). (n=3 for the rats number of each group)

### 7. Inflammation in balloon-inflated brains was also influenced by balloon deflation time

On the 3rd day after the operation, the brains were collected, and the expression of proinflammatory cytokines, including IL-1β, IL-6 and TNF-α, was assessed using ELISA kits. Consistent with Piezo1 expression, the levels of all three cytokines in rats in the10-s deflation time group were significantly higher than those in the 1-min deflation time group (P<0.05). However, the levels of IL-1β, IL-6 and TNF-α were not significantly different between the groups with 5-min and 1-min deflation times (P>0.05) **(Fig. 7)**.

**Figure 7.**
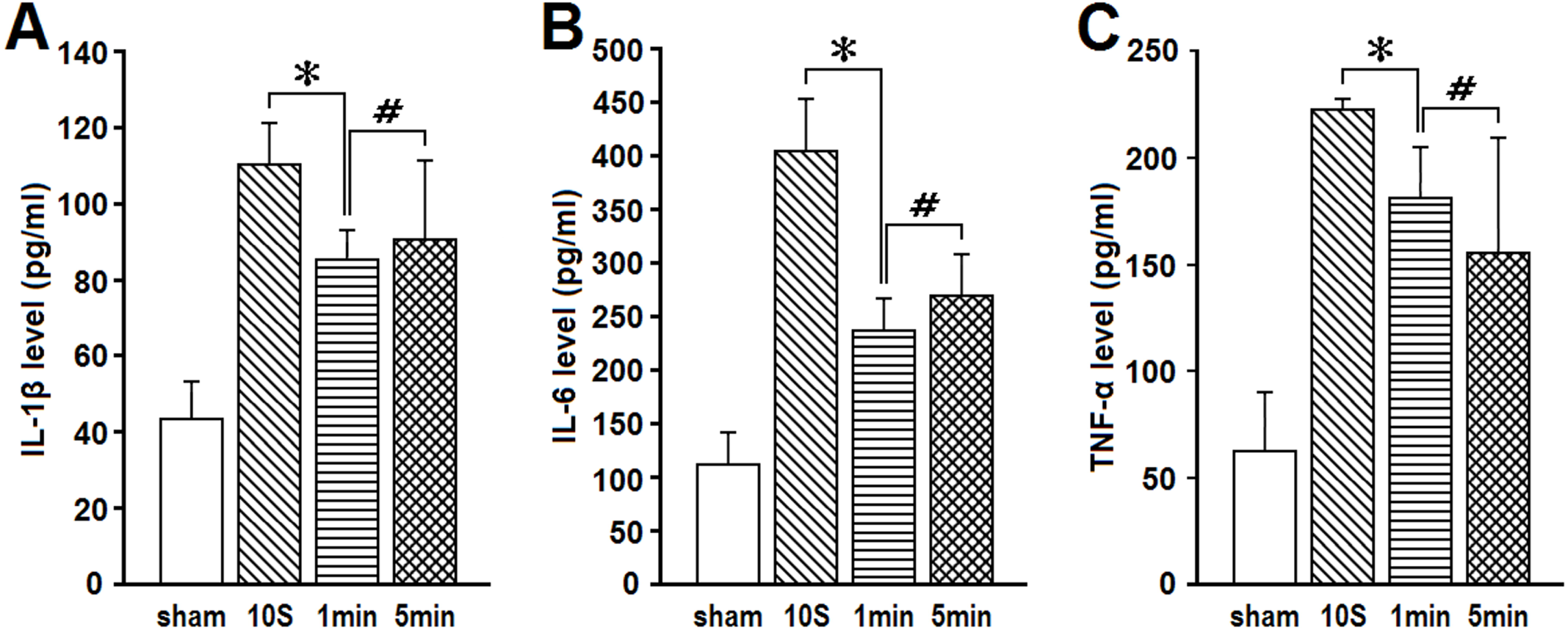
Comparison of levels of proinflammatory cytokines among groups with different balloon deflation times. Levels of classic proinflammatory cytokines IL-1β, IL-6 and TNF-α were quantitatively analyzed and compared among different groups. “sham” means the balloon was inserted but not inflated. “10 s”, “1 min”, and “5 min” refers to the deflation time after the balloon was inflated for 30 min. Student’s t test was used for statistical analysis. Asterisks indicate significant differences (P<0.05), and hashes indicate no prominent difference (P>0.05). (n=3 for the rats number of each group)

## Discussion

In the present study, we first established a stable balloon-inflated rat model to mimic pure mechanical compression-induced brain injury after ICH and demonstrated that the compression induced abnormal behavior in rats. Second, we found that the mechanosensitive ion channel Piezo1 was expressed in compressed brain tissue, and the expression level was prominently dependent on the existence of compression. Third, the levels of three proinflammatory cytokines (IL-1β, IL-6, TNF-α) were higher in balloon-inflated rat brains than in sham control brains. Moreover, after balloon inflation, the deflation velocity played an active role in compression injury in model rats. Finally, GsMTx4, an antagonist of Piezo1, improved animal behavior and reduced proinflammatory cytokine expression in model rats.

The balloon inflation brain injury model, mimicking only the mass effects and no blood components of hematoma, emerged around the end of the 1980s (Sinar *et al*., 1987) but was not widely used and was soon discarded in favor of other animal models of ICH such as auto-blood injection and collagenase injection (MacLellan *et al*., 2008; Manaenko *et al*., 2011). These two models well mimic the biochemical changes of ICH and have greatly improved our knowledge of injury from thrombin, hemoglobin, heme, iron ions, edema, inflammation, and oxidative stress excitotoxicity (Keep *et al*., 2012; Zhou *et al*., 2014). Meanwhile, the mass effect has largely been ignored. This bottleneck may be partially attributed to the absence of a bridge that links “mass compression” and “injury pathology”, thereby limiting further explanation for why a mechanical stimulus results in tissue injury. Considered as “true mechanosensitive ion channels in mammals”, the Piezo family has attracted much attention since it was first identified (Coste *et al*., 2010; Martinac and Poole, 2018). Having been studied in various regions, it was reminiscent of the ICH injury. In ischemia models, the artery occlusion time is usually 90 min (Liu and McCullough, 2011); therefore, in our pilot study, rats were exposed to balloon inflation for 30 min or 1 h to avoid the influence of potential ischemia injury. However, mortality associated with a 1-h inflation time was too high to maintain robust survival after modeling. Therefore, we considered 30 min to be the best balloon inflation time for the establishment of a stable model.

The complicated nature of brain injury after ICH onset makes dividing the damage into “primary” and “secondary” injury difficult. In recent year, studies on SAH (subarachnoid hemorrhage) have used the term “early brain injury” (EBI) to refer to injury that occurs early in the process, usually within 72 h. Animal studies have shown that neuroinflammation and neuronal apoptosis are two major processes associated with EBI, of which the detailed mechanisms need to be determined (Ji and Chen, 2016). Analogically considering, we chose the 3rd day after model establishment as the observation point.

In the present study, we found that pure mechanical compression can result in hemiplegia symptoms in rats and that Piezo1 was expressed in the brain tissue surrounding the balloon inflation focus, while the rest of the brain tissue was not positive for Piezo1 immunofluorescence. A similar phenomenon was observed in studies on the expression of the TRPA1 channel (one subtype of TRP channels), which indicated that TRPA1 mRNA and protein expression levels were very low in physiological conditions and only upregulated with tissue damage (Moran *et al*., 2004; Barritt and Rychkov, 2005). Interestingly, Piezo1 mRNA expression has been shown to be increased after bladder obstruction in mice, indicating that the expression of Piezo1, as a mechanosensitive channel, is influenced by mechanical force (Michishita *et al*., 2016). Another report supports it by indicating that Piezo1 expression pattern is affected by mechanical stress (Blumenthal et al., 2014). Together, these results suggest that Piezo1 is only expressed in brain tissue upon tissue compression. Another possibility is that Piezo1 expression is maintained at a very low level not detectable using immunohistochemistry and is then prominently enhanced upon compression by an exogenous mass. Anyway, our results indicate that Piezo1 expresses in brain tissue and that compression by exogenous mass results in a prominent increase in its expression level.

In our study, Piezo1 expression was accompanied by an increase in proinflammatory cytokine expression. Calcium signaling might act as a mediator, as the chief downstream event of Piezo1 activation has been shown to be Ca^2+^ influx (Bagriantsev *et al*., 2014).Early studies have indicated that Ca^2+^ dysregulation constitutes the basis of neuronal injury, as many factors involved in neuronal functions are associated with Ca^2+^ signaling (Aarts and Tymianski, 2005). Among several calcium-related pathways, Janus kinase (JAK)-signal transducer and activator of transcription proteins (STAT) 1/3, a classic cytomembrane signaling pathway that can be mechanically activated (Pan *et al*., 1999), was recently detected to modulate cytokines, including interferons and interleukins (Vignali and Kuchroo, 2012; Boisson-Dupuis *et al*., 2012; Chmielewski *et al*., 2014), suggesting that the role of this pathway in mechanotransduction should be further investigated. These reports, together with our findings, indirectly suggest that the mediator between Piezo1 and neuroinflammation is Ca^2+^ influx, serving as the early cause of neuronal injury; thus, more research will be needed in the future.

GsMTx4 has been reported to be an efficient antagonist of Piezo1; however, GsMTx4 may be more accurately defined as a negative modulator of Piezo1 as its inhibitory effect can be overcome by stronger mechanical stimuli (Gnanasambandam *et al*., 2017). The target of GsMTx4 is mainly closed Piezo1 rather than opened Piezo1, although the existence of other binding states cannot be excluded. GsMTx4 has been reported to have no effect on the whole-cell current of resting cells, indirectly supporting the hypothesis that Piezo1 is inactive in quiescent conditions (Bae *et al*., 2011). In the present study, GsMTx4 was shown to improve the behavioral outcome and attenuate neuroinflammation without prominently affecting Piezo1 expression. Our data might suggest that Piezo1 was inhibited by GsMTx4 treatment functionally rather than structurally. In addition, GsMTx4 has been shown to inhibit the mechanosensitive current and volume decrease in rat kidney fibroblast NRK49 cells, indicating that Piezo1 might be the sensor of cell swelling in this system (Hua *et al*., 2010). Notably, cytomembranes are stretched when cells swell, which may be one mechanism by which mechanosensitive channels are activated (Ranade *et al*., 2015).

We exposed the rats to different balloon deflation times and obtained interesting results. After the balloon was inflated for 30 min, deflation of the balloon within 10s led to higher Piezo1 expression, higher proinflammatory cytokine expression and worse functional behavior than deflation within 1 min or 5 min. These results indicate that compression removal is not always beneficial but rather depends on the velocity of deflation. This phenomenon may be attributed to the structure and function of Piezo1, which has already been partially and tentatively illuminated: the sensitivity of Piezo1 to mechanical stimuli is not limited to compression but rather the “change in cytomembrane tension” (Lewis and Grandl, 2015; Cox *et al*., 2016; Zhao *et al*., 2016; Wu *et al*., 2017). Therefore, Piezo1 may be speculated to be activated by the deformation of the cytomembrane, which occurs during both compression and decompression. Thus, we deduce that in ICH, although clot evacuation may eliminate hematoma damage, the resulting stretching of the cytomembrane may result in new injury, the extent of which depends on the velocity of this process: the severer, the worse; the milder, the better. This phenomenon that both compression and decompression in ICH brains activates Piezo1 could be summarized as “bidirectional”. Hence, it could partially explain the occurrence of a poor functional outcome after hematoma evacuation despite good imaging results (Aiyagari, 2015).

Importantly, our results do not indicate hematoma evacuation is unnecessary but rather suggest that there may be some previously unrecognized harmful effects accompanying the benefits of hematoma removal that might explain the disconnect between successful hematoma removal and poor outcome. However, further research is necessary to confirm this hypothesis.

Prokaryotic MSCs have been recognized in *Escherichia coli* for approximately 30 years, but the exploration of MSCs in animals has been difficult (Kung *et al*., 2010; Zhao *et al*., 2016). Before Piezos were discovered, there were only three well-identified MSCs in the animal kingdom: ENaC/DEG, TRP and K2P (Chalfie, 2009), and the evidence of their function in mammals was still lacking (Delmas and Coste, 2013). Most identified MSCs showed ability of mechanotransduction but could not be mechanical gated or represented mechano-gating modulation but possessed no mechanosensitivity, indicating that they are not “real” mechanical signaling channels (Medhurst *et al*., 2001; Talley *et al*., 2001; Chalfie, 2009). Currently, the roles of DEG/EnaC and TRP channels in the mechanotransduction of mammals are still unclear. The novel Piezo channel family was identified in 2010 and attracted the interest of researchers (Coste *et al*., 2010). TRP and DEG/EnaC channels are widely accepted to be essential for mechanotransduction in invertebrates, but mechanotransduction in mammals likely depends more on Piezo proteins (Ranade *et al*., 2015). The relationship between genetic mutations of *PIEZO1*/*PIEZO2* and human diseases has further proven their importance (Albuisson *et al*., 2013). At present, at least 25 mutations of Piezo1 have been identified to be responsible for different human diseases (Kung *et al*., 2010), and at least 12 mutations of Piezo2 have been associated with arthrogryposis (McMillin *et al*., 2014; Okubo *et al*., 2015), among which two have been electrophysiologically identified to be damaging to channel inactivity, leading to a prominent increase in Ca^2+^ influx (Coste *et al*., 2013). Up to now, the study on Piezo1 is proceeding to Piezo2, whether Piezo2 is structurally or functionally similar to Piezo1 remains unclear. Thus, Piezo2 and Piezo1 and their respective roles in the nervous system may differ. For example, many approaches that were able to activate Piezo1 were unable to activate Piezo2 (Syeda *et al*., 2015). Furthermore, a stretching stimulus was only able to activate a small current through Piezo2 channel (success rate of 50% for Piezo2 and 90% for Piezo1) (Lee *et al*., 2014; Coste *et al*., 2015). The two homologous isomers may have different activation mechanisms: Piezo1 is a polymodal sensor of various mechanical stimuli, while the sensing activity of Piezo2 may be narrower (Kung *et al*., 2010).

## Conclusion

The secondary injury associated with ICH has traditionally been thought to begin at clot lysis and blood component release, which occur approximately 72h after attack onset. In the present study it indicated that there was inflammation-like response within 30 minutes after formation of hematoma, which might be the super early injury mediated by Piezo1, with the potential mechanism of Ca^2+^ influx resulting neuroinflammation. In addition, our study showed that the improvement resulting from mechanical compression removal depends on the removal velocity, which may be attributed to the “bidirectional” response of Piezo1 to mechanical stimuli.

We have suggested that the activation and upregulation of Piezo1 could be the upstream and early event of ICH-induced brain injury, establishing the first *in vivo* evidence for the role of Piezo1 in blood-brain neuroinflammation after ICH. Although these findings cannot be immediately translated to the clinic, our study provides new insight into the mechanism of the mass effect after ICH and might lead to the identification of new treatment targets, which will require further research.

## Author contributions

Yichi Zhang designed and carried out the model establishment, immunohistochemistry and molecular biology experiments, analyzed the results of the studies above, and wrote the paper. Gang Wang carried out behavioral test, took part in model establishment, molecular biology experiments and data analysis. Lifei Lian took part in data collection, discussing the results and revising the manuscript. Yongjie Xiong and Feng Xu took part in the interpretation of the results and discussion. Guo Li coordinated the statistic analysis. Zhouping Tang and Furong Wang gave helpful advices to the design of experiments. Minjie Xie helped with improvement of the whole experiment protocol, quantity controlling of all experiments and the manuscript revising. Suiqiang Zhu conceived and designed the whole study, supervised all of the experiments, and revised the whole manuscript.

## Declaration

The authors declare no conflict of interest.

## Abbreviations

ICH: Intracerebral hemorrhage
MSCs: Mechanical sensitive ion channels
TRP: transient receptor potential channels
K2P: Double pore K+ channels
DRG: dorsal root ganglion
SPF: Specific pathogen free
EBI: early brain injury

